# Intranasal hemagglutinin protein boosters induce robust mucosal immunity and cross-protection against influenza A viral challenge

**DOI:** 10.1101/2024.10.10.616291

**Authors:** Miyu Moriyama, Gisele Rodrigues, Jiping Wang, Andrew Hudak, Huiping Dong, Robert J. Homer, Drew Weissman, Shuangge Ma, Akiko Iwasaki

## Abstract

Licensed parenteral influenza vaccines induce systemic antibody responses and alleviate disease severity but do not efficiently prevent viral entry and transmission due to the lack of local mucosal immune responses. Here, we describe intranasal booster strategy with unadjuvanted recombinant hemagglutinin (HA) following initial mRNA-LNP vaccination, Prime and HA. This regimen establishes highly protective HA-specific mucosal immune memory responses in the respiratory tract. Intranasal HA boosters provided significantly reduced viral replication compared to parenteral mRNA-LNP boosters in both young and old mice. Correlation analysis revealed that slightly increased levels of nasal IgA are significantly associated with a reduced viral burden in the upper respiratory tract. Intranasal boosting with an antigenically distinct H1 HA conferred sterilizing immunity against heterologous H1N1 virus challenge. Additionally, a heterosubtypic intranasal H5 HA booster elicited cross-reactive mucosal humoral responses. Our work illustrates the potential of a nasal HA protein booster as a needle- and adjuvant-free strategy to prevent infection and disease from influenza A viruses.

**One Sentence Summary:** Adjuvant-free nasal booster induces protective immunity against influenza infection.

## INTRODUCTION

Seasonal influenza epidemics present a significant public health challenge, leading to approximately half a million deaths globally each year (*1*). Despite vaccines being the most effective preventive measure against seasonal influenza, their effectiveness remains limited, ranging between 19% and 54% (Centers for Disease Control and Prevention. Seasonal influenza vaccine effectiveness studies, 2014–2024. https://www.cdc.gov/flu/professionals/vaccination/effectiveness-studies.htm<u>. Accessed 13 July 2024</u>). While these influenza vaccines can reduce the complications from seasonal influenza, they do not efficiently prevent human-to-human viral transmission. This limitation is due to the inability of these parenteral vaccines to elicit mucosal immune responses including IgA and tissue-resident memory cells in the respiratory mucosa, which are crucial for rapid anamnestic responses that block the spread of respiratory viruses (*2, 3*). The development of vaccines capable of inducing such protective local immune responses holds promise for controlling the transmission of influenza and other respiratory viruses (*4*).

Recent studies have shown that vaccine regimens combining parenteral priming with intranasal boosting can induce robust mucosal immune responses and enhance protection against viral challenges, as demonstrated in SARS-CoV-2 animal models (*5, 6*). In a previous study, we introduced the “Prime and Spike” strategy, in which intramuscular priming with a Spike-encoding mRNA-LNP vaccine (prime) was followed by intranasal boosting with an unadjuvanted, unformulated, trimeric Spike protein (Spike). This approach generated strong protective mucosal immune responses and reduced viral transmission (*6*). Currently, multiple clinical trials are underway to test the efficacy of influenza mRNA vaccines (*7*).

Building on these findings, this report explores the use of the HA mRNA-LNP vaccine platform for priming and demonstrates that intranasal unadjuvanted HA protein boosters, termed “Prime and HA”, can evoke sterilizing immunity against respiratory viral challenge. Our study highlights the potential of the “Prime and HA” strategy to enhance influenza vaccine effectiveness and contribute to better control of seasonal influenza epidemics and future pandemics.

## RESULTS

### Intranasal unadjuvanted HA protein boosters confer near-sterilizing immunity in mice primed with intramuscular mRNA-LNP vaccination

To evaluate the protective efficacy of the parenteral priming followed by intranasal boosting strategy against lethal influenza virus infection, we immunized Balb/c mice intramuscularly (IM) with an mRNA-LNP vaccine encoding influenza A/PR8 HA and boosted them with intranasal (IN) unadjuvanted recombinant HA protein (Fig. 1A). A sub-protective dose (0.05μg) of mRNA-LNP for IM priming protected 20% of the mice from death and did not prevent severe weight loss (Fig. 1B-D) and a high viral load in both the upper and lower respiratory tracts (Fig. 1E-H). A single IN HA protein booster (Prime and HA; P+HA) substantially mitigated weight loss (Fig. 1C-D) but was still insufficient to fully protect mice from death (Fig. 1B) and viral replication (Fig. 1E-H). In contrast, an IN HA protein booster following two doses of IM mRNA-LNP (Prime and boost and HA; P+B+HA) or P+HA (Prime and HA and HA; P+HA+HA) completely protected mice from morbidity and mortality (Fig. 1B-D) and minimized lung tissue pathology (Fig. 1I-J) to levels comparable to those with three doses of mRNA-LNP (Prime and boost and boost; P+B+B). Notably, IN HA protein booster (P+B+HA) conferred near-sterilizing immunity, as evidenced by the absence of infectious virus in the nasal wash and bronchoalveolar lavage fluid (BALF) (Fig. 1E-F) and only minimal viral RNA remnants in the lungs (Fig. 1G-H), a level of protection not achieved by the parenteral mRNA-LNP booster (P+B+B). Additionally, animals that received P+HA+HA displayed reduced viral replication compared with P+B+B (Fig. 1E-H), highlighting the robust boosting effect of IN HA protein administration following mRNA-LNP priming.

**Fig. 1.**
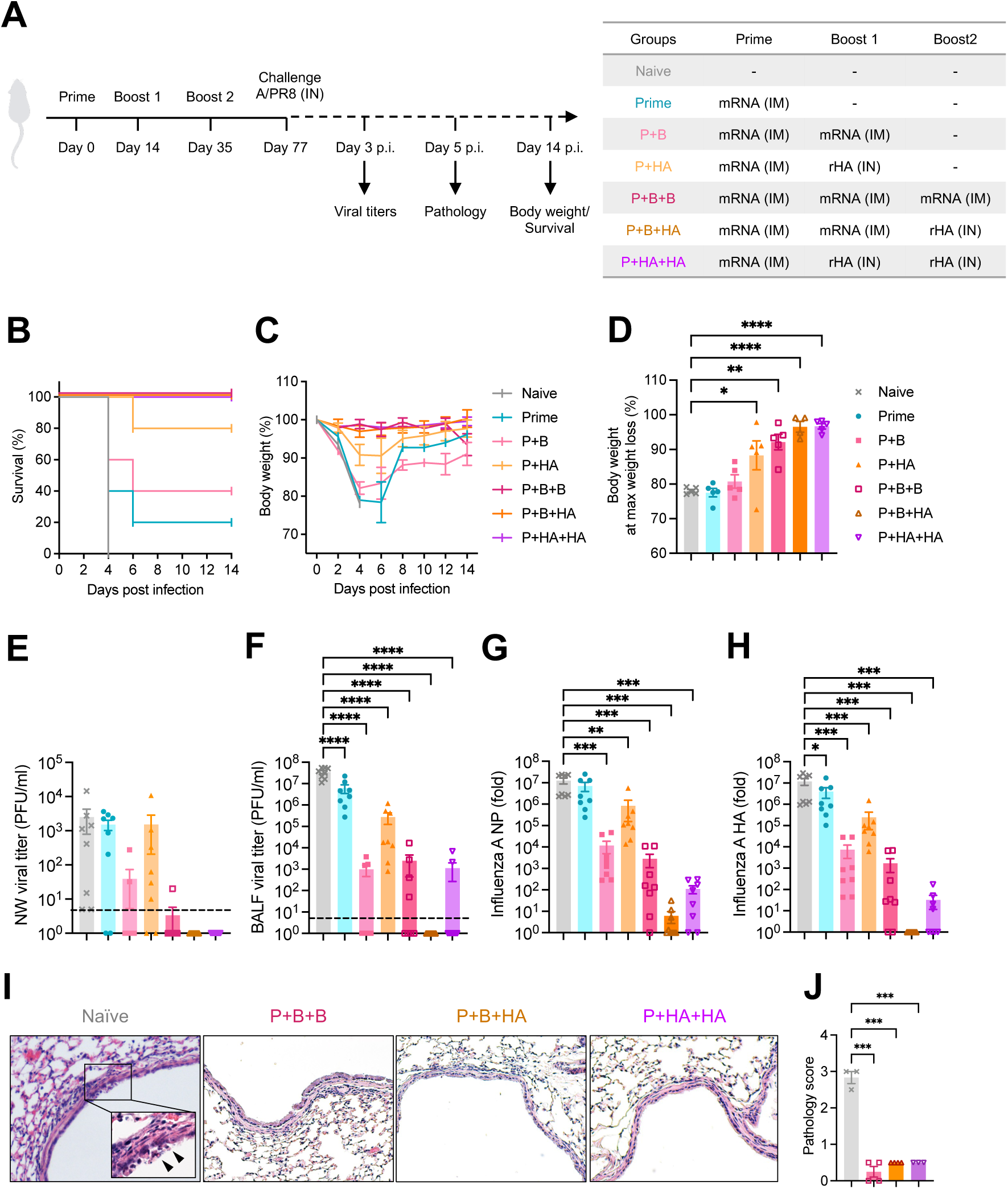
Intranasal unadjuvanted HA protein boosters following parenteral mRNA-LNP immunization provide robust protection. (**A**) Balb/c mice were immunized with 0.05μg of PR8 HA mRNA-LNP intramuscularly and 5μg recombinant PR8 HA protein intranasally as indicated. Six weeks after the third dose, mice were challenged with 10^4^ PFU of A/PR8. (**B** to **D**) Survival and body weight were monitored for 14 days. (**E** to **H**) Nasal wash, BALF, and lung tissue were collected 3 days after infection, and infectious viral titers in nasal wash (**E**) and BALF (**F**) were determined by plaque assay. Viral RNA levels in the lung tissue were determined by RT-qPCR. (**I** and **J**) The immunized Balb/c mice were challenged with 10^5^ PFU of A/PR8. Lung tissue was collected 5 days after infection for pathological assessment by H&E staining. Morphologic cell death and degeneration are observed in the naive lung (arrowheads in the inset). Data are representative or pooled from at least two independent experiments. Error bars are shown in mean+/- S.E.M. *, p<0.05; **, p< 0.01; ***, p< 0.001; ****, p< 0.0001

We next assessed the efficacy of the vaccine strategy against a different influenza virus strain over an extended period post-immunization. The protective immunity was robustly induced following P+B+HA and P+HA+HA against another H1N1 strain, A/Michigan/45/2015 (MI15), and was sustained for at least 12 weeks after the second boost (Fig. S1A-E). Conversely, a single IM immunization with a quadrivalent influenza vaccine (QIV) containing MI15 HA, the current standard of care, only partially protected mice from weight loss and did not substantially reduce the viral load at this time point (Fig. S1A-E). Further, IN MI15 HA booster given to QIV-primed mice failed to confer antiviral resistance. These results underscore the superior ability of IN HA protein boosters to block infection in the nose and lung over parenteral mRNA-LNP boosters following the initial series of IM mRNA-LNP vaccination, but not after QIV.

### P+B+HA and P+HA+HA establish mucosal immune memory responses

Parenteral delivery of mRNA-LNP vaccine induces robust circulating humoral and cellular immune responses (*8*), but not resident memory CD4+ and CD8+ T cells or secretory IgA in the respiratory tract (*9, 10*). Given that P+B+HA and P+HA+HA regimens reduced viral shedding compared to P+B+B, we hypothesized that IN HA boosters elicit robust local immune responses in the respiratory mucosa, the primary site of influenza virus entry and replication. To determine whether the intranasal delivery of recombinant HA protein following two doses of parenteral mRNA-LNP vaccination establishes protective mucosal immune memory, we assessed HA-specific humoral and T cell responses 6 weeks post-final booster, when memory lymphocyte populations are established. Consistent with the previous findings, P+B+B led to the induction of systemic (Fig. 2A) and local (Fig. 2B) IgG responses but failed to induce either systemic or local mucosal IgA responses (Fig. 2C-E). By contrast, P+B+HA and P+HA+HA resulted in robust mucosal IgA responses in the upper and lower respiratory tracts (Fig. 2D&E) in addition to systemic and local IgG responses (Fig. 2A&B). We further examined tissue-resident memory T cell (TRM) responses in the lung and found a significant increase in polyclonal CD4+ TRMs and antigen-specific CD8+ TRMs in the P+B+HA and P+HA+HA groups (Fig. 2F&G). Conversely, P+B+B induced CD8+ TRMs but did not stimulate polyclonal CD4+ TRM responses (Fig. 2F&G). These results demonstrated the unique capacity of nasal HA protein boosters to establish robust local mucosal immune memory responses.

**Fig. 2.**
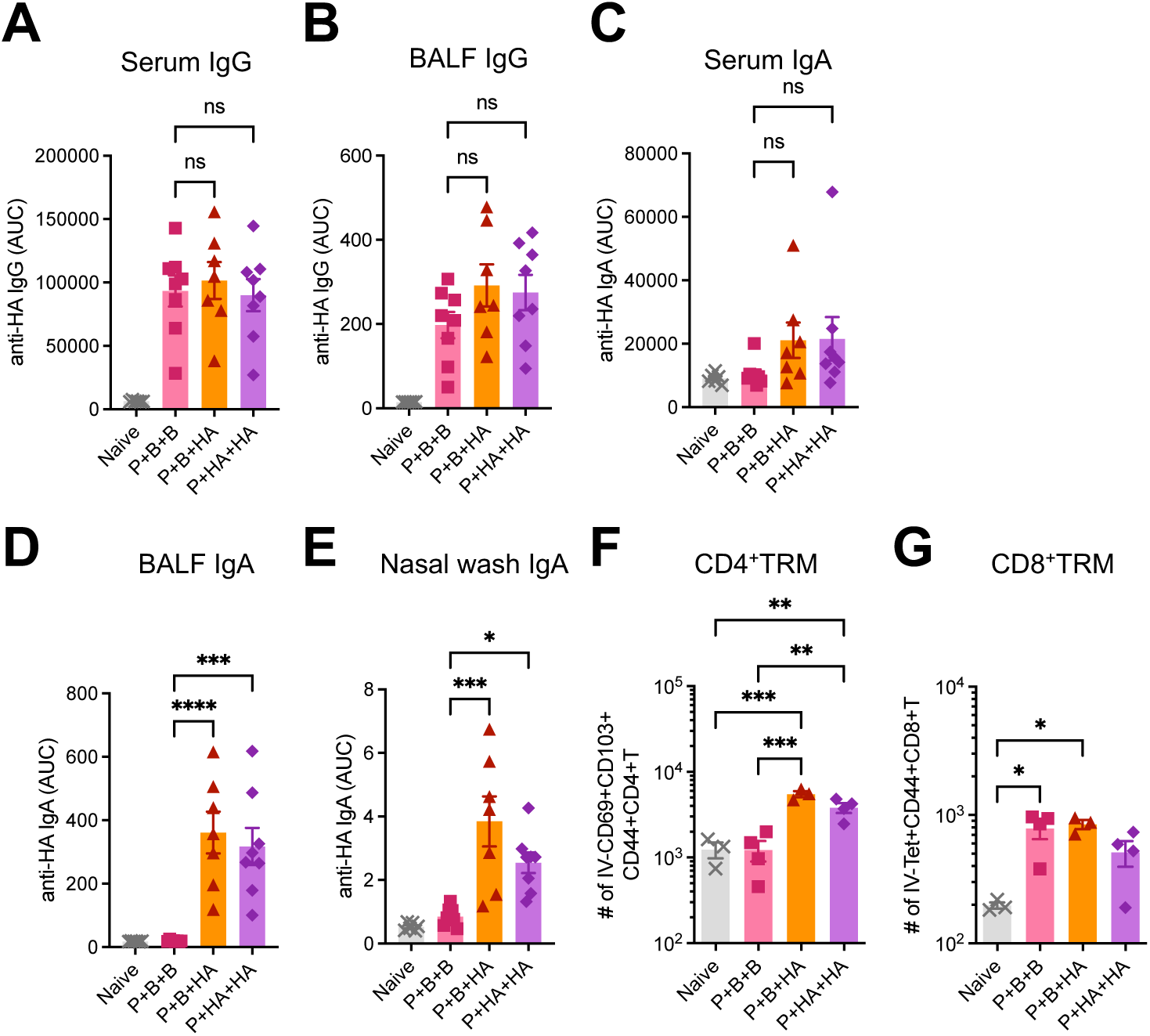
Intranasal HA boosters following mRNA-LNP immunizations establish mucosal humoral and cellular immune responses. Balb/c mice were primed and boosted with 0.05μg of PR8 HA mRNA-LNP intramuscularly and received either an additional dose of PR8 HA mRNA-LNP (Prime and boost and boost; P+B+B) or 5μg recombinant PR8 HA protein intranasally (Prime and boost and HA; P+B+HA). Another group of Balb/c mice were primed with 0.05μg of PR8 HA mRNA-LNP intramuscularly and received two doses of 5μg recombinant PR8 HA booster intranasally (Prime and HA and HA; P+HA+HA). Six weeks after the second boost, serum, BALF, nasal wash, and lung tissue were corrected and assessed for humoral (**A** to **E**) and cellular (**F** and **G**) immune responses. (**A** to **E**) PR8 HA-specific systemic and mucosal IgG and IgA responses were measured by ELISA. (**F** and **G**) Polyclonal CD4+ resident memory T cells (TRMs) and HA-specific CD8+ TRMs in lung tissue were quantified by flow cytometry. Data are representative or pooled from two independent experiments. Error bars are shown in mean+/- S.E.M. *, p<0.05; **, p< 0.01; ***, p< 0.001; ****, p< 0.0001

### IgG and IgA are strong mucosal immune correlates of protection

Systemic HA-binding antibody titers correlate well with resistance to influenza and have long served as a standard for measuring vaccine effectiveness (*11*). However, to advance the development of next-generation influenza vaccines, a deeper understanding of the correlates of protection, particularly mucosal correlates, is crucial (*4, 12, 13*). To identify the mucosal correlates of protection against influenza virus infection, we measured HA-specific IgA and IgG levels in the nasal wash and BALF from the same cohort as Figure 1D-G. Spearman correlation coefficients were calculated between the levels of mucosal antibodies and protection, as reflected by viral shedding after the challenge. We focused on the groups that received IN HA boosters (P+HA, P+B+HA, and P+HA+HA) for this correlation analysis, as the groups without IN HA boosting lacked mucosal IgA responses (Fig. S2). Our analysis revealed strong and significant negative correlations between BALF HA-specific IgG and both BALF viral titers (r=-0.7373, p<0.0001) and lung influenza NP RNA (r=-0.813, p<0.0001), as well as lung influenza HA RNA (r=-0.8419, p<0.0001) (Fig. 3A, bottom row). These negative correlations were as strong as the correlations between serum HA-specific IgG and viral load (Fig. 3C). Further, we observed similarly strong negative correlations between BALF HA-specific IgA levels and viral shedding in the BALF (Fig. 3A, top row).

**Fig. 3.**
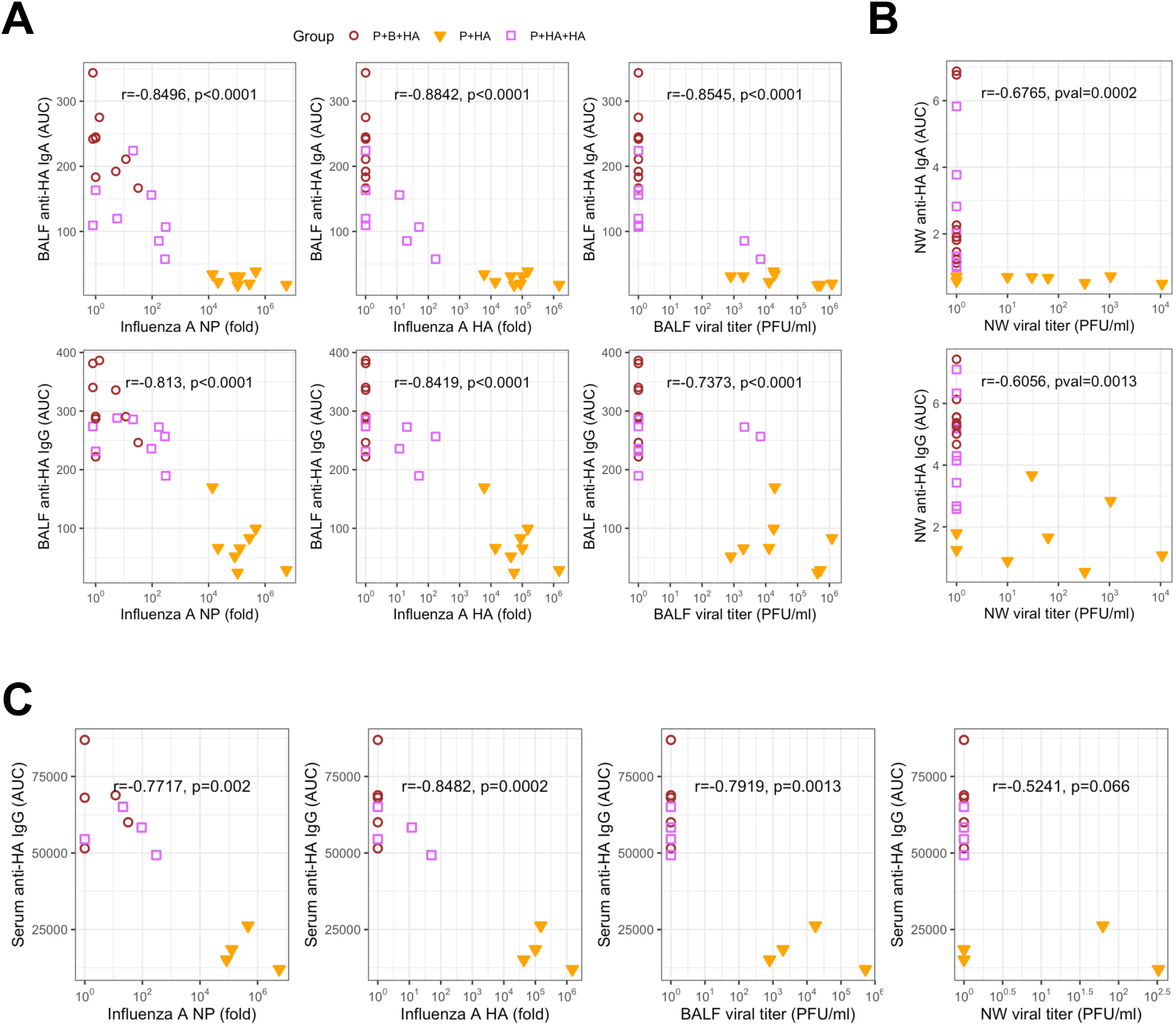
Mucosal IgA and IgG inversely correlate with viral burden. (**A** to **C**) Balb/c mice were immunized with one or two doses of 0.05μg of PR8 HA mRNA-LNP intramuscularly and boosted with 5μg recombinant PR8 HA protein intranasally (P+HA or P+B+HA). Another group of Balb/c mice were primed with 0.05μg of PR8 HA mRNA-LNP intramuscularly and received two doses of 5μg recombinant PR8 HA booster intranasally (Prime and HA and HA; P+HA+HA). Six weeks after the second boost, mice were challenged with 10^4^ PFU of A/PR8. Nasal wash, BALF, and lung tissue were collected 3 days after infection. Infectious viral titers, viral RNA levels, and PR8 HA-specific antibody responses were determined. Spearman correlations denoted by R between anti-HA antibody levels and either viral titers or viral RNA levels in BALF (**A**), nasal wash (**B**), and serum (**C**).

Additionally, we found significant negative correlations between nasal wash HA- specific IgG and nasal viral titers (Fig. 3B). Notably, even extremely low levels of anti-HA IgA were sufficient to completely block viral replication in the nasal wash (Fig. 3B, top row). These findings demonstrate the protective roles of mucosal IgG and IgA, in addition to serum IgG, in upper and lower respiratory tracts.

### Intranasal HA boosters provide protective immune responses in old mice

Advanced age is a significant risk factor for influenza virus and other respiratory infections (*14*). Despite the critical need for protection in this population, the immunogenicity and effectiveness of seasonal influenza vaccines among adults older than 65 years of age are lower than those observed in younger populations (*15*). To assess whether our vaccination strategy offers enhanced protection in aged hosts, we compared the protective efficacy of the parenteral mRNA-LNP vaccination regimen (P+B+B) and intranasal HA boosting regimens (P+B+HA or P+HA+HA) in older mice. Previous studies have shown that both resistance to respiratory viral infection (*16*) and mRNA-LNP-induced antibody responses (*17*) are reduced in 1-year-old mice compared to younger mice. Female Balb/c mice older than 14 months were immunized with P+B+B, P+B+HA, or P+HA+HA regimens, and their protective efficacy was evaluated after 6 weeks by monitoring body weight, survival, and viral burden after challenge (Fig. 4A). All vaccinated mice, except for one out of seven immunized with P+HA+HA, survived and were protected from weight loss following an otherwise 100% lethal infection (Fig. 4B-C). Although all immunization groups significantly reduced viral shedding in BALF (Fig. 4D), a considerable viral burden remained (Fig. 4D, F) in older mice. Yet, P+B+HA reduced BALF viral titers by a thousand-fold, offering more efficient protection than P+B+B (Fig. 4D). In contrast, the same immunization regimens provided a substantial reduction in viral burden and conferred sterilizing immunity in young mice (Fig 4E, G), illustrating the attenuated vaccine-induced viral resistance in aged hosts. We hypothesized that the superior reduction in respiratory viral load observed with P+B+HA was associated with stronger mucosal antibody responses. Indeed, P+B+HA elicited higher systemic and mucosal HA-specific IgG and IgA responses compared with P+B+B (Fig. 4H-K). These data indicate that the mucosal HA booster provides better immune responses than mRNA-LNP booster, correlating with a significantly lower viral replication in aged mice.

**Fig. 4.**
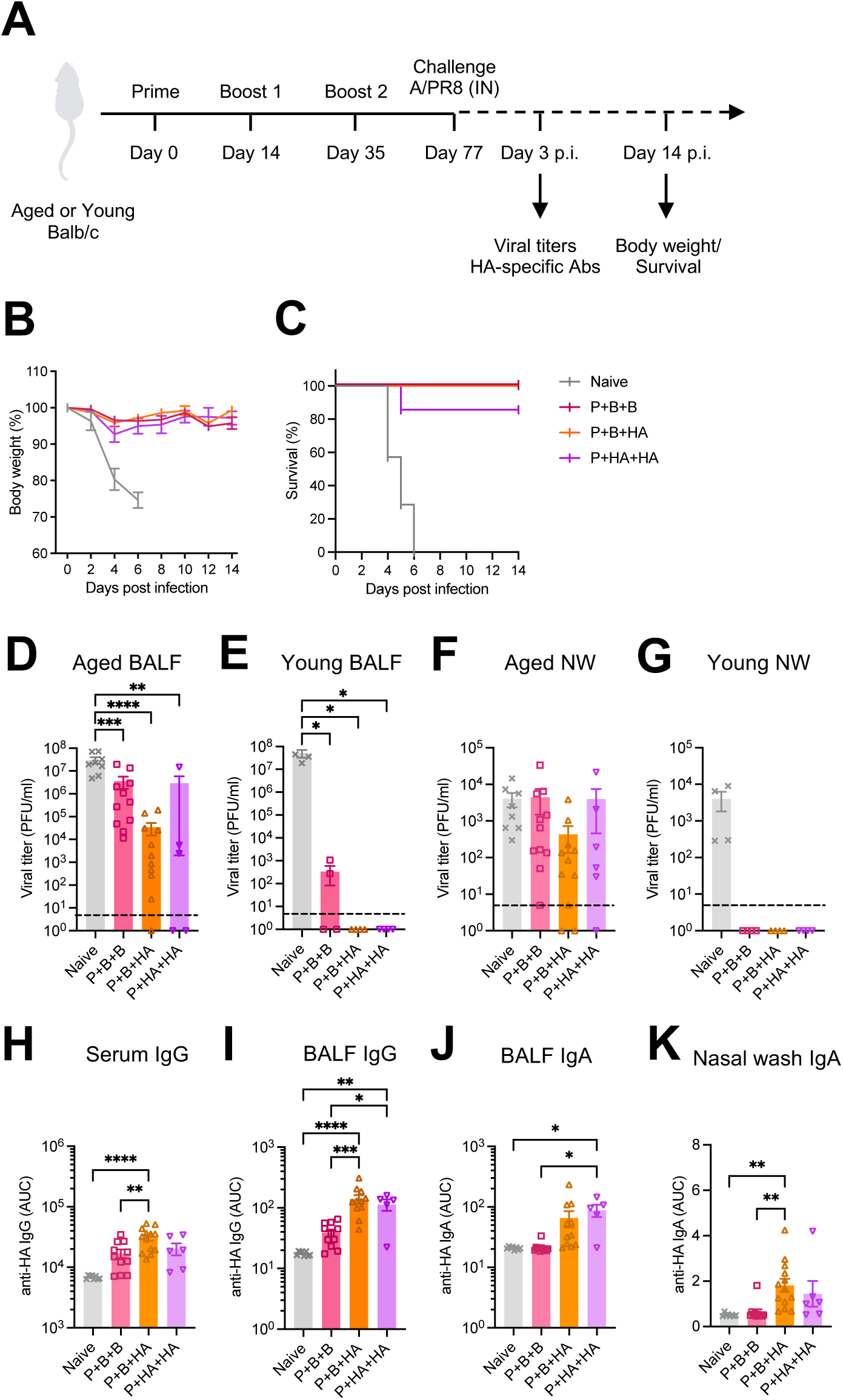
Intranasal HA administration boosts protective immunity in aged mice. (**A**) Aged Balb/c mice were primed and boosted with 0.05μg of PR8 HA mRNA-LNP intramuscularly and received either an additional dose of PR8 HA mRNA-LNP (Prime and boost and boost; P+B+B) or 5μg recombinant PR8 HA protein intranasally (Prime and boost and HA; P+B+HA). Another group of aged mice were primed with 0.05μg of PR8 HA mRNA-LNP intramuscularly and received two doses of 5μg recombinant PR8 HA booster intranasally (Prime and HA and HA; P+HA+HA). A paired young cohort was prepared for the viral load assessment. Six weeks after the second boost, mice were challenged with 10^4^ PFU of A/PR8. Serum, nasal wash, and BALF were collected 3 days after infection. (**B** and **C**) Body weight and survival were monitored for 14 days after challenge. (**D** to **G**) Viral load in BALF (**D** and **E**) and nasal wash (**F** and **G**) were determined by plaque assay. (**H** to **K**) PR8 HA-specific systemic and mucosal IgG and IgA responses were measured by ELISA. Data are pooled from at least two independent experiments. Error bars are shown in mean+/- S.E.M. *, p<0.05; **, p< 0.01; ***, p< 0.001; ****, p< 0.0001

### Intranasal heterosubtypic H5 HA boosters generate cross-reactive humoral immunity

Next, we investigated whether the intranasal boosting with H5 HA protein could induce cross-reactive mucosal humoral responses in the hosts previously exposed to an H1N1 viral infection. Balb/c mice who were infected with a sublethal dose of A/California/04/09 (Cal09) strain 12 months prior were intranasally boosted with H5 HA protein twice (Fig. 5A). The prior infection with Cal09 alone did not result in significant cross-reactive antibody responses against H5 HA, with only a moderate increase in H5-specific IgG in the BALF (Fig. 5B-E). After two doses of intranasal H5 booster, there was a marked elevation in both systemic (Fig. 5B) and local (Fig. 5C) H5-specific IgG responses, as well as local H5-specific IgA responses (Fig. 5D-E). These findings demonstrate the ability of intranasal heterosubtypic H5 HA protein boosters to induce cross-reactive humoral immune responses, extending beyond the influenza virus subtype previously encountered.

**Fig. 5.**
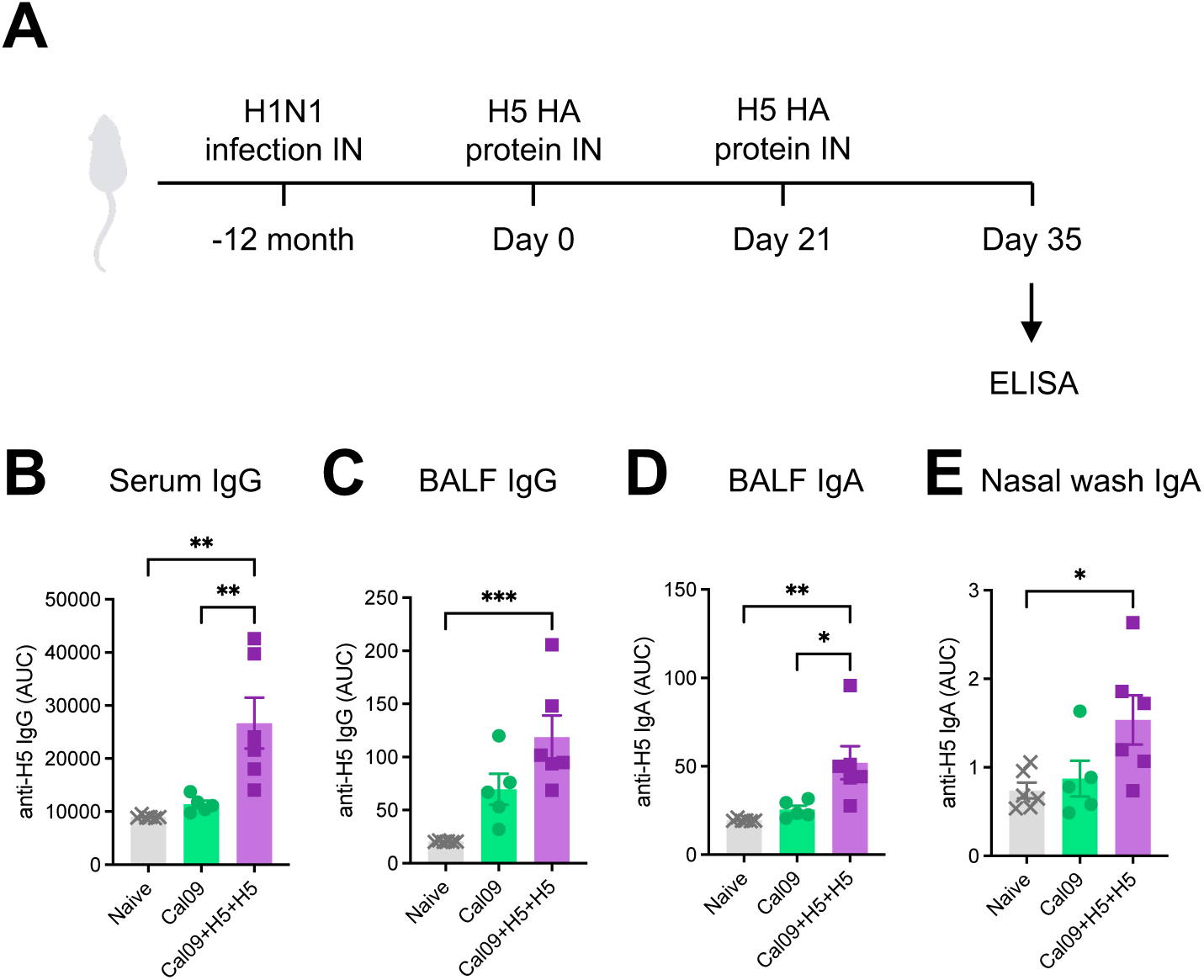
Intranasal heterosubtypic H5 HA boosters elicit H5-reacting mucosal immune responses. (**A**) Balb/c mice infected with a sublethal dose (10^5^ PFU) of A/California/04/09 (H1N1) 12 months prior were immunized twice with 4μg recombinant H5 (VN04; A/Vietnam/1194/2004) HA protein intranasally (Cal09+H5+H5) or remained untreated (Cal09). Two weeks after the second boost, serum, nasal wash, and BALF were collected and analyzed for VN04-reacting antibody responses. (**B** to **E**) VN04 HA-specific systemic and mucosal IgG and IgA responses were measured by ELISA. Data are representative of two similar experiments. Error bars are shown in mean+/- S.E.M. *, p<0.05; **, p< 0.01; ***, p< 0.001

### Intranasal heterologous H1 HA boosters elicit cross-protective immunity

The antigenic evolution of seasonal influenza viruses substantially decreases the efficacy of seasonal influenza vaccines from one year to the next (*12*). We sought to assess the potential of intranasal heterologous H1 HA booster to generate protective immune responses against an antigenically distant strain. To this end, mice were primed with mRNA-LNP vaccine encoding PR8 HA and then intranasally boosted twice with a recombinant HA cocktail containing a 1:1 mixture of PR8 HA and antigenically distant MI15 HA protein, which differ in HA amino acid sequence by 18.2% (Prime and HA mix and HA mix; P+HAm+HAm). As a control, another group received three doses of PR8 HA mRNA-LNP (P+B+B) (Fig. 6A). Six weeks after the final dose, the immunized mice were challenged with an MI15 strain and assessed for protection based on weight loss, viral load, and cross-reacting antibody responses against MI15. The mice immunized with P+HAm+HAm exhibited only slight weight loss on day 2 post-infection but quickly recovered (Fig. 6B). In addition, there was no detectable infectious viral load in either the nasal wash (Fig. 6C) or the BALF (Fig. 6D) of P+HAm+HAm mice, indicating that the intranasal heterologous HA boosters elicited robust cross-protective immunity against MI15. Consistently, significant induction of systemic and mucosal MI15-specific IgG (Fig. 6E-F) and mucosal IgA (Fig. 6G-H) was observed in the P+HAm+HAm group. By contrast, mice immunized with P+B+B lost body weight to levels similar to the Naïve group (Fig. 6B). P+B+B did not alter the viral titers in the nasal wash (Fig. 6C) but reduced those in the BALF by half (Fig. 6D) of Naive. This reduction in BALF viral titer was associated with the specific elevation of MI15-reacting IgG responses in the BALF (Fig. 6F), indicating the partial protective capacity of cross-reactive BALF IgG. These data further suggest the requirement of other immune responses including mucosal IgA and systemic IgG for full protection. Collectively, these data demonstrate that intranasal heterologous HA boosting confers sterilizing immunity against an antigenically distant strain within the H1N1 subtype (Fig. 6C-D).

**Fig. 6.**
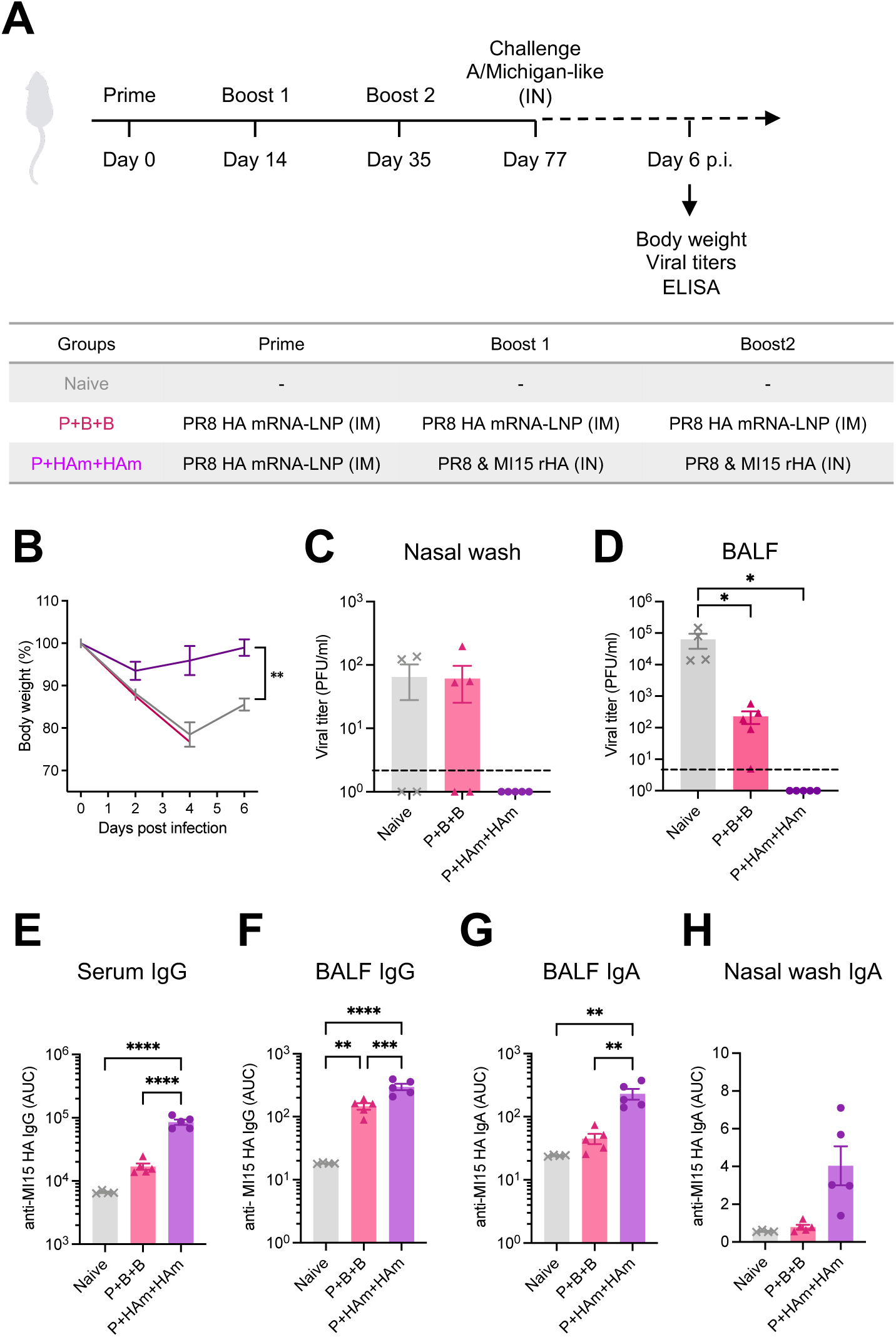
Heterologous intranasal HA boosters expand the breadth of protective immunity against the heterologous H1N1 strain. (**A**) Balb/c mice were primed with 0.05μg of PR8 HA mRNA-LNP intramuscularly and boosted twice with recombinant HA cocktail (2.5μg each of PR8 and MI15 HA) intranasally (Prime and HA mix and HA mix; P+HAm+HAm) or received three doses of PR8 HA mRNA-LNP intramuscularly (Prime and boost and boost; P+B+B). Six weeks after the second boost, mice were challenged with 5000 PFU of A/Michigan/45/2015. Body weight was monitored for 6 days (**B**). On day 6, serum, nasal wash, and BALF were collected. (**C** and **D**) Viral load in nasal wash (**C**) and BALF (**D**) were determined by plaque assay. (**E** to **H**) MI15 HA-specific systemic and mucosal IgG and IgA responses were measured by ELISA. Data are representative of two similar independent experiments. Error bars are shown in mean+/- S.E.M. Statistical significance was tested using two-way ANOVA for Figure 3B. *, p<0.05; **, p< 0.01; ***, p< 0.001; ****, p< 0.0001

## DISCUSSION

In this study, we demonstrate the ability of IN HA protein boosters following IM HA-encoding mRNA-LNP priming to elicit robust protective mucosal immune responses and confer sterilizing immunity against influenza virus infection. While IN HA booster and IM mRNA-LNP booster similarly induced systemic and local HA-specific IgG responses, mucosal anti-HA IgA antibodies were uniquely induced by the IN HA booster. Protection conferred by Prime and HA is long-lasting and was demonstrated even in older mice. By using heterologous HA boosters, we were able to demonstrate cross-reactive antibodies against the newly introduced HA in the mucosa and sterilizing immunity following heterologous viral challenge. The simplicity of needle-free administration emphasizes the potential of IN HA booster as an alternative strategy to reduce disease severity. Our findings also validate that the previously established “Prime and Spike” approach (*6*) can be effectively applied to different pathogens. Crucially, Prime and HA provides a simple vaccine approach to establish resistance against infection, thus inhibiting onward transmission of influenza virus.

Here, we used the mRNA-LNP platform to generate pre-existing systemic immunity, which allowed an intranasal adjuvant-free recombinant HA protein booster to induce potent respiratory mucosal immune responses. Even though an IN HA booster moderately promoted the recovery of QIV-primed mice, its booster effect was significantly more pronounced in mRNA-LNP-primed mice. Previous research has shown that a single intradermal mRNA-LNP injection generates robust splenic single and polyfunctional CD4+ and follicular helper T cells (TFH) in mice, unlike IM immunization with inactivated influenza virus (*18–20*). The variability in the booster effect likely reflects differences in the quality and magnitude of immune responses induced by different priming vaccines.

We demonstrated that IM prime-boost immunization of HA encoding mRNA-LNP followed by an IN recombinant HA booster, which we term P+B+HA, elicits robust mucosal immune responses. The P+B+HA regimen not only prevented disease progression and lung pathology but also provided sterilizing immunity against a lethal influenza virus challenge, a protection not afforded by parenteral mRNA-LNP vaccination control (P+B+B). The enhanced protection with P+B+HA was accompanied by robust mucosal IgA responses in upper and lower respiratory tracts, suggesting the substantial role of mucosal IgA antibodies in limiting viral replication. The importance of local IgA antibodies is further supported by their significant negative Spearman correlations with viral shedding in BALF (Fig. 3A) and the inverse relationship between nasal IgA levels and nasal viral titer (Fig. 3B). These results align with previous studies illustrating the importance of nasal secretory IgA in reducing viral shedding in humans (*21, 22*) and underscore the importance of local mucosal immunity as the first line of defense in the respiratory mucosa. Recent studies on COVID-19 vaccines have shown that mucosal vaccines delivered via inhalation or intratracheal routes can induce near-sterilizing protective immunity in non-human primates and humans (*23, 24*). For instance, Ye et al. reported that their nanoparticle-based inhaled vaccine prevents aerosol transmission of SARS-CoV-2 in hamsters (*24*). Future research should explore whether our nasal booster approach can efficiently prevent viral transmission.

Developing alternative vaccine strategies to improve protection for the older adults who are at higher risk of developing severe disease is a key focus for next-generation influenza vaccines. Successful vaccination of older adults is a benchmark for novel vaccine approaches, given their typically reduced responsiveness compared to younger populations (*13*). To enhance the immunogenicity of seasonal influenza vaccines in older adults, MF-59 adjuvanted or high-dose formulations have been approved (*25, 26*). In this study, we employed the potent mRNA-LNP platform for priming and assessed the protective efficacy of IN HA booster versus standard IM mRNA-LNP booster in aged mice. Consistent with findings in young mice, P+B+HA mounted higher systemic and local antibody responses and significantly reduced viral burden compared with the standard P+B+B control in older hosts with weakened immune systems. Given the engagement of distinct immune cell populations and signaling pathways, the boosting effect of IN HA and IM mRNA-LNP can be differently impacted by aging. While mRNA-LNP vaccines elicit multitude of innate sensors for adaptive immune activation (*27–29*), aging-related degradation of TRAF3 and diminished IFN expression may hinder the adjuvanticity of the vaccines (*30*). Nevertheless, for transmission-blocking vaccine strategy, our data show that IN HA protein booster is still effective in reducing viral load in older mice.

Limitations of our study include the use of inbred mice, which may not reflect the immunology or physiology of human respiratory tract. Second is the lack of demonstration of transmission-blocking capabilities of the Prime and HA approach. Since mice are not naturally capable of transmission (*31*), future studies involving established models of transmission such as ferrets are needed to address this question.

One of the major challenges for influenza vaccines is the emergence of immune escape mutants due to the rapid antigenic evolution of the viral surface glycoproteins. In this study, we addressed this challenge by demonstrating two sequential doses of IN heterologous or heterosubtypic HA boosters to expand the breadth of antibody reactivity in mice previously vaccinated or infected. Administering an antigenically diverged H1 HA protein as an IN booster following priming with an original HA-encoding mRNA-LNP vaccine elicited cross-protective mucosal immunity. Given the potential pandemic threat posed by the recently emerged bovine H5N1 influenza (*32*), we also explored the potential of IN H5 HA booster to establish cross-reactive immunity in hosts with prior exposure to an H1N1 virus. Our findings with the heterosubtypic H5 HA IN booster doses were sufficient to induce cross-reactive mucosal IgG and IgA responses in mice that recovered from an H1N1 virus one year prior. The promise of the “Prime and HA” to confer transmission- and infection-blocking immunity in humans merits further investigation.

## MATERIALS AND METHODS

### Study Design

The aim of this study was to evaluate the efficacy of an intranasal unadjuvanted HA protein booster following intramuscular mRNA-LNP priming. This included a comprehensive virological and immunological assessment, such as the detection of HA-specific resident memory T cells using H-2K(d) tetramers. We employed a well-established mouse model of influenza virus infection, utilizing Balb/c mice and the mouse-adapted A/Puerto Rico/8/34 (H1N1) strain. A/Michigan/45/2015 (H1N1) strain was used to evaluate heterologous protection.

### Mice

Six to ten-week-old female Balb/c mice were purchased from Charles River Laboratories or bred in-house. Fourteen to Twenty-month-old female Balb/c mice were used as aged mice. All animal experiments in this study complied with federal and institutional policies of the Yale Animal Care and Use Committee.

### Cells and viruses

Madin-Darby canine kidney (MDCK) cells were maintained in MEM supplemented with 1% Penicillin-Streptomycin and 10% heat-inactivated fetal bovine serum (FBS) at 37°C. A/Puerto Rico/8/34 (H1N1) was kindly provided by Dr. Hideki Hasegawa (National Institute of Infectious Diseases, Japan). A/California/04/09 (H1N1) was kindly provided by Dr. Adolfo Garcia-Sastre (Icahn School of Medicine at Mount Sinai). A/Michigan/45/2015 (H1N1) was kindly provided by Dr. Michael Schotsaert (Icahn School of Medicine at Mount Sinai). Virus stocks were propagated in allantoic cavities from 10- to 11-day-old fertile chicken eggs for 2 days at 35°C. Viral titers were determined by standard plaque assay procedure. Briefly, serial 10-fold dilutions of nasal wash and BALF were prepared in PBS containing 0.1% bovine serum albumin (BSA). Aliquots of 200μl of diluted samples were inoculated into a confluent monolayer of MDCK cells in 6-well plates and incubated for 1 hour at 37°C. After incubation, cells were washed with PBS and overlaid with 2ml agar-containing MEM supplemented with TPCK-treated trypsin. Forty-eight hours later, cells were stained with crystal violet and the number of plaques in each well was counted. All experiments using live influenza virus were performed in biosafety level 2 laboratories and animal facilities with approval from the Yale Environmental Health and Safety office.

### Vaccines

Nucleoside-modified, influenza-HA encoding mRNA-LNPs were produced as described previously (*29*). Briefly, nucleoside-modified mRNAs were produced by using N1-methylpseudouridine (m1Ψ)-5′-triphosphate from codon-optimized A/Puerto Rico/8/1934 or A/Michigan/45/2015 HA-encoding mRNA production plasmids. The mRNAs were designed to contain 101 nucleotide-long poly(A) tails and 5’ capping. Cellulose-purified, nucleoside-modified mRNAs were encapsulated in lipid nano particles (LNPs) by self-assembly process and stored at −80°C until use. Fluzone quadrivalent influenza vaccine 2017-2018 (QIV) was obtained from BEI resources. Recombinant A/Puerto Rico/8/34 HA (Cat#11684-V08H1), A/Michigan/45/2015 HA (Cat#40567-V08H1), and A/Vietnam/1194/2004 (Cat#11062-V08H1) were purchased from Sino Biological.

### Vaccination and infection

For intramuscular immunization, mice were anesthetized by isoflurane inhalation and injected with 0.05μg mRNA-LNP or 25μl QIV adjusted to 50μl in PBS. For intranasal vaccination or infection, mice were fully anesthetized by intraperitoneal injection of ketamine and xylazine, and intranasally inoculated with 50μl of PBS containing recombinant HA protein or influenza virus.

### Measurement of anti-HA antibodies

Influenza HA-reacting antibodies were detected by enzyme-linked immunosorbent assay (ELISA) as previously described (*33*). Briefly, 96-well MaxiSorp plates were coated with 2μg/ml of HA protein in carbonate buffer overnight at 4°C and blocked with PBS containing 4% FBS for 1 hour at room temperature. Serum, nasal wash, and BALF samples were diluted in PBS containing 4% FBS and applied to the plates. Following overnight incubation at 4°C, plates were washed with 0.1% Tween-20 containing PBS and incubated with horseradish peroxidase (HRP)-conjugated anti-mouse IgG or IgA.

After 1 hour of incubation at room temperature, plates were washed with 0.1% Tween-20 containing PBS. TMB substrate (eBioscience) was added to each well and 1N sulfuric acid was added to terminate the reaction. The absorbance of 450 nm wavelength was recorded.

### Intravenous labeling and lung cell isolation

Circulating lymphocytes were labeled by the previously described intravenous labeling method (*6*). Briefly, the APC-Cy7-labeled anti-mouse CD45 antibody (Cat#103116, BioLegend) was diluted in PBS and injected intravenously 3 minutes before the tissue harvest. Lungs were harvested and processed as previously described (*9*). Briefly, lungs were minced with scissors and digested in RPMI1640 media containing 1mg/ml collagenase A, 30μg/ml DNase I at 37°C for 45 min. Digested lungs were then filtered through a 70μm cell strainer and treated with ACK buffer for 2 min. After washing with PBS, cells were resuspended in PBS with 1% FBS and subjected to surface marker staining.

### Flow cytometry

Cells were blocked with mouse Fc Block in the presence of Live/Dead Fixable Aqua (Cat# L34966, 1:1000, Thermo Fisher) for 20 min at 4°C. Staining antibodies were added and incubated for 30 min at 4°C. The stained cells were washed with 2mM EDTA-PBS and resuspended in 100μl 2% PFA for 15 min at 4°C. After fixation, cells were washed and resuspended in PBS with 1% FBS and analyzed on Attune NxT (Thermo Fisher). The data obtained were analyzed using FlowJo software (Tree Star). Staining antibodies are as follows (PerCP-Cy5.5 anti-mouse CD8α (Cat#103057, 1:400, BioLegend), PE-Cy7 anti-mouse CD69 (Cat#104512, 1:200, BioLegend), AF700 anti-mouse CD4 (Cat#100536, 1:200, BioLegend), BV421 anti-mouse CD103 (Cat#121422, 1:200, BioLegend), BV605 anti-mouse TCRβ (Cat# 109241, 1:200, BioLegend), BV711 anti-mouse CD44 (Cat# 103057, 1:200, BioLegend)). PE-conjugated H-2K(d) tetramer (Influenza A HA 533-541 IYSTVASSL) was provided by the NIH Tetramer Core Facility.

### Quantitative PCR

Mouse lung tissues were minced with scissors and lysed with TRIzol reagent (Invitrogen). Total RNA was extracted using an RNeasy mini kit (QIAGEN) and reverse transcribed into cDNA using an iScript cDNA synthesis kit (Bio-Rad). RT-PCR was performed by CFX96 Touch Real-Time PCR detection system (Bio-Rad) using iTaq SYBR premix (Bio-Rad) and the following primers (5’-3’): Influenza NP (Forward: AGAACATCTGACATGAGGAC, Reverse: GTCAAAGGAAGGCACGATC), influenza HA (Forward: AGTGCCCAAAATACGTCAGG, Reverse: GGCAATGGCTCCAAATAGAC).

### Pathological assessment

Lung tissue was fixed in 4% paraformaldehyde overnight and transferred into 70% ethanol. Paraffin embedding, sectioning, and hematoxylin and eosin (H&E) staining of the tissue were performed by Yale Pathology Tissue Services. Pathological findings were identified and scored by a pulmonary pathologist through blinded sample evaluation using the following criteria: Airway injury is identified based on a composite of the evaluation of morphologic cell death and degeneration and inflammation including acute inflammation using the percent and extent of the circumference involved. Immune cell infiltration is determined based on 1, minimum visible infiltration; and 4, circumferential massive infiltration of lymphocytes and eosinophils. Notably, in all cases, the extent of inflammation was relatively low and the score was primarily driven by epithelial injury.

### Statistical analysis

Statistical significance was tested using one-way analysis of variance (ANOVA) with Tukey’s multiple comparison test unless otherwise stated. A two-way ANOVA test or a two-tailed t-test were used where indicated. For correlation analysis, Spearman rank correlation coefficients were calculated to capture all monotonic relationships. No missing data were present in the cohort that received IN HA boosters and no obvious outliers were observed upon visual inspection of the data distribution. Statistical significance of correlations was assessed using two-tailed t-test. P-values of <0.05 were considered statistically significant. Correlation analyses were performed using R (version 4.4.1). Prism 9 were used for all other statistical analyses.

## Acknowledgments

We thank Melissa Linehan and Pavlina Baevova for their technical and logistical assistance. We thank Michael Schotsaert for kindly providing A/Michigan/45/2015 strain. We thank the NIH Tetramer Core Facility for providing H-2K(d) tetramer. We thank the BEI resources repository for providing Fluzone QIV. We thank Tianyang Mao for the critical reading of the manuscript. The graphic was created by BioRender.

## Author contributions

M.M. and A.I. conceived of and designed the project. M.M., G.R., A.H., H.D., and R.J.H. performed experiments. M.M., J.W., S.M., and A.I. analyzed and interpreted data. D.W. supplied reagents. D.W., S.M., and A.I. supervised the study. M.M. and A.I. wrote the original manuscript and all authors reviewed and provided feedback on the manuscript.

## Data availability

All study data are included in the main text or Supplementary materials.

## Supplementary Figures

**Fig. S1.**
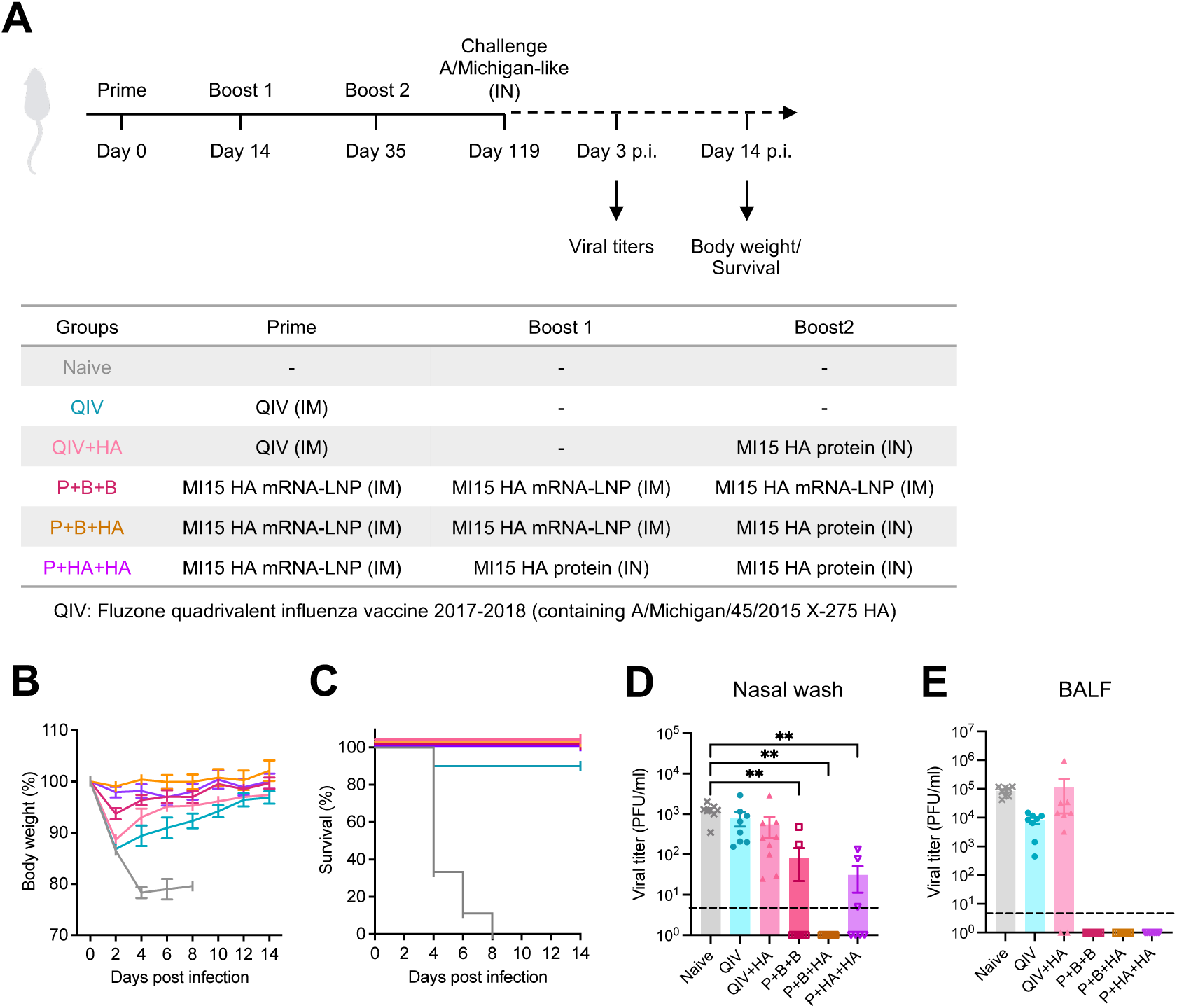
Protective efficacy of intranasal HA boosters sustains for at least 12 weeks. **(A)** Balb/c mice were immunized with 0.05μg of A/Michigan/45/2015 (MI15) HA mRNA-LNP intramuscularly and 5μg recombinant MI15 HA protein intranasally as indicated. Quadrivalent influenza vaccine (QIV) served as a standard of care control. Twelve weeks after the second boost, mice were challenged with 10^4^ PFU of A/Michigan/45/2015. (**B** and **C**) Body weight and survival were monitored for 14 days after the challenge. (**D** and **E**) Nasal wash and BALF were collected 3 days after infection, and infectious viral titers in the nasal wash (**D**) and BALF (**E**) were determined by plaque assay. Data are pooled from two independent experiments. Error bars are shown in mean+/- S.E.M. *, p<0.05;

**Fig. S2.**
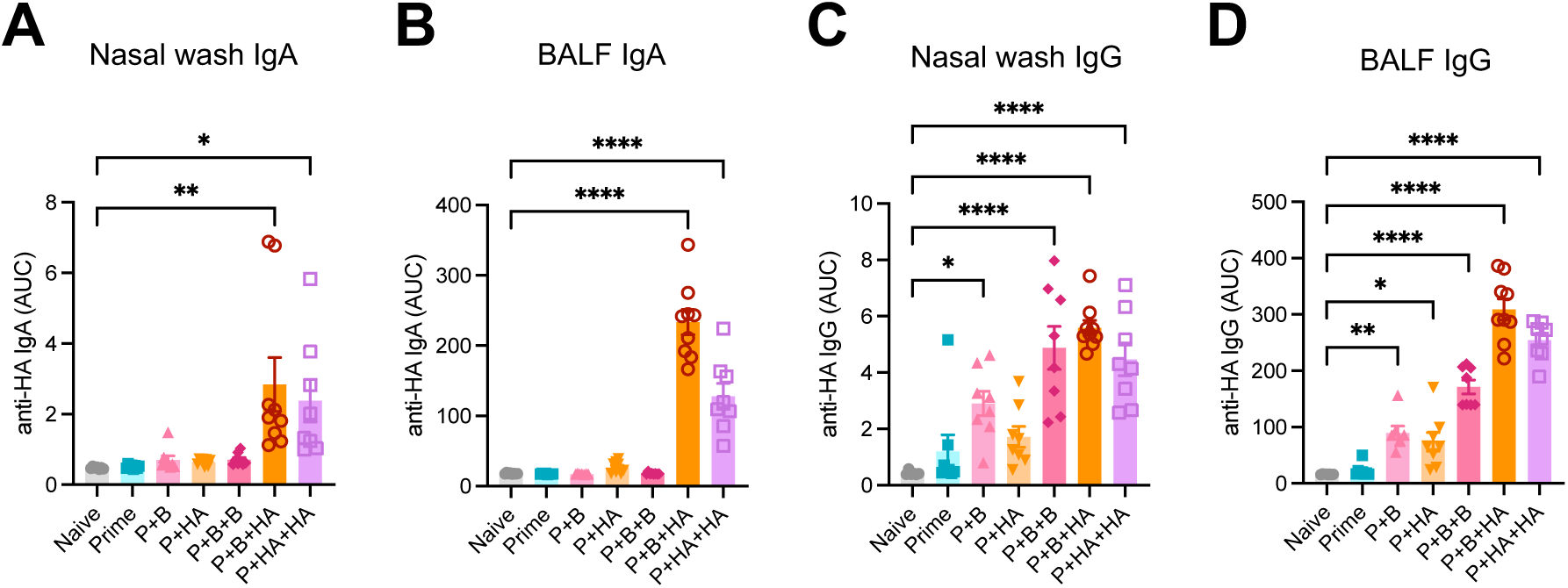
HA-specific humoral immune responses. (**A** to **D**) Balb/c mice were immunized with 0.05μg of PR8 HA mRNA-LNP intramuscularly and 5μg recombinant PR8 HA protein intranasally as indicated in Figure 1A. Six weeks after the second boost, mice were challenged with 10^4^ PFU of A/PR8. Three days after infection, nasal wash and BALF were collected and subjected to the measurement of PR8 HA-specific IgG and IgA responses by ELISA. Data are pooled from two independent experiments. Error bars are shown in mean+/- S.E.M. *, p<0.05; **, p< 0.01; ****, p< 0.0001

